# Complement C3 interacts with cytochrome *c* to influence myocardial apoptosis during heart ischemia/reperfusion

**DOI:** 10.1101/2023.03.15.532779

**Authors:** Zhou Fang, Haekyung Lee, Junying Liu, Karen A. Wong, Lewis M. Brown, Ming Zhang

## Abstract

Myocardial ischemia/reperfusion (I/R) elicits an acute inflammatory response involving complement factors. Previous animal studies showed that circulation complement C3 was deposited in the ischemic myocardium flooded with oxygenated blood upon reperfusion. Recently, we reported that myocardial necrosis was decreased in C3^−/−^ mice after heart I/R. The current study used in the same heart model to test the effect of C3 on myocardial apoptosis. Our results showed that myocardial apoptosis was increased in C3^−/−^ mice after heart I/R. Further, comparative proteomics analyses found that cytochrome *c* was present in the myocardial C3-complex following I/R. These results indicate that C3 can interact with cytochrome *c* in the cytosol of cardiomyocytes during myocardial I/R, which may sequester cytochrome *c* and thus reduce the number of cells undergoing apoptosis. In summary, our findings raise the possibility of a new mechanism affecting cell death relevant to pathologic conditions such as ischemia: a circulating innate immune factor, i.e. complement, can interact with intracellular factor(s), and influence the types of cell death that occur.

## 1. Introduction

Myocardial ischemia/reperfusion (I/R) elicits an acute inflammatory response involving complement factors of the innate immune system [1–3]. Complement inhibitors, which have the potential to limit inflammation, showed promises in preclinical studies [4–8]. However, limited positive results were obtained with the few inhibitors studied in clinical trials of myocardial I/R injury [3]. In particular, anti-complement C5 (Pexelizumab) failed to meet the primary endpoints in cardiac patients who had undergone coronary artery bypass graft surgery [9] or percutaneous transluminal coronary intervention [10]. One explanation of the conflicting clinical trial results is that they targeted downstream components in the complement pathway, e.g., C5, leaving earlier activators such as C3 unaffected. Activation of the earlier complement factors would have affected pathways such as those leading to cell death with the potential to influence the outcome.

Previous animal studies by us and others showed that under pathological conditions such as I/R injury, circulation C3 was deposited in the ischemic myocardium flooded with oxygenated blood upon reperfusion [11,12]. Recently we reported that in the mouse heart model, myocardial necrosis was decreased in C3^−/−^ mice [13]. This implied that in WT mice during I/R, C3 acts to promote necrosis. Besides necrosis, basic research has indicated that apoptosis plays an important role in cardiac I/R injury [14-16]. Whether C3 is involved in apoptosis during myocardial I/R is unclear.

## 2. Materials and Methods

### 2.1. Mouse model of myocardial I/R injury

Complement C3 knockout (C3 ^−/−^) mice and WT (C57BL/6) mouse strains were obtained from the Jackson Laboratory (Bar Harbor, ME) and maintained at the SUNY Downstate Medical Center Department of Laboratory Animal Resources. Genotyping was provided by GeneTyper (New York, NY). Male mice were used at 10-12 weeks of age (weight 26-30 g) in accordance with the requirements of the NIH and the Institutional Animal Care and Use Committee (IACUC) of SUNY Downstate Medical Center. The protocol was approved by the IACUC of SUNY Downstate Medical Center (Approval #11-10276).

We employed an established model of myocardial I/R injury model [12,17]. Mice were anesthetized using sodium pentobarbital (60 mg/kg, i.p.), intubated and ventilated with a mouse ventilator (Harvard Apparatus, MA). Following sternotomy, the left anterior descending artery (LAD) was ligated for 1 h; occlusion of the LAD was confirmed by the appropriate color change of myocardial tissue and the ST elevation on ECG; reperfusion was verified by the reversed color change of the left ventricle and the appropriate ECG changes. Postoperative management included fluid replacement with normal saline and pain relief with the analgesic buprenorphine (0.1 mg/kg, intramuscularly). The mice were sacrificed the hearts were harvested at 3 hrs of reperfusion for proteomic study and 24 hrs of reperfusion for histopathologic analyses.

### 2.2. Apoptosis analyses of mouse heart undergoing I/R

TUNEL Assay: Frozen sections were cut from the heart slices and stained using a TACS 2TdT TUNEL kit according to the manufacturer’s directions (Trevigen, Gaithersburg, MD). TUNEL-positive cells in sections from all 4 slices of each heart were quantified and expressed as per mm^2^ of the cryosection and the average was obtained for the 3 mice of each group.

Caspase-3 activation: cryosections were stained with a rabbit Ab to activated caspase-3 (Cell Signaling Technology, MA) using published procedures [18,19], and then with a secondary mouse anti-rabbit IgG tagged with FITC (Millipore, MA). Positively stained areas were quantified and expressed as positive area (μm^2^)/total area (mm^2^).

### 2.3. Comparative proteomics

Comparative proteomics with label-free shotgun profiling was carried out at Quantitative Proteomics Center, Columbia University. Proteins in heart samples were solubilized using Rapigest (Waters Corp., MA), and digested with trypsin. Peptides were separated in a 120-min LC run on a 0.75-μm ID x 25-cm reverse phase 1.7 μm particle diameter C18 column and analyzed on a Synapt Q-TOF mass spectrometer. MS and MS^E^ spectra were analyzed using an MS^E^/Identity^E^ algorithm. In addition, the Elucidator Protein Expression Data Analysis System software (Rosetta Biosoftware, MA), which can import Identity^E^ data, were used for advanced multivariate statistical analysis, false discovery control, principal components analysis, multidimensional scaling, and cluster analysis. To identify the C3-specific signaling network in necrosis, proteins identified from proteomics were mapped to existing biological networks by Ingenuity Pathway Analysis (QIAGEN, CA)

### 2.4. ELISA measurement of cytochrome c in the C3-immunocomplex

A sandwich ELISA was used to verify the presence of cytochrome *c* in the C3-complex. The wells of a 96 well plate was coated with an anti-C3 Ab (Complement Technology, TX) to capture the C3-complex in the heart cytosolic fractions. After washing to remove unbound factors and blocking with BSA, an anti-cytochrome *c* Ab (Enzo, NY) was used to detect the presence of cytochrome *c*.

### 2.5. Western blotting of cytochrome c

Cytosolic fractions were isolated using a published method [20]. Briefly, hearts were homogenized in a Dounce homogenizer in cytosolic extraction buffer. Lysates were centrifuged (600 g, 10 min, 4 °C) and the supernatant collected and further centrifuged (10,000 g, 10 min, 4 °C). The cytosolic extract (supernatant) was collected. The protein (20 μg) from each cytosolic extract was analyzed by Western blotting following 12% SDS-PAGE. Membranes were probed with a sheep anti-mouse cytochrome *c* Ab (Enzo, NY) followed by a donkey anti-sheep IgG coupled to horse radish peroxidase (R&D, MN) and developed with an ECL Western blotting kit (Thermo Scientific, NJ). The intensities of the cytochrome *c* bands were normalized to that of constitutively expressed β-actin and expressed as relative intensity. Quantification of bands of interest was carried out using the ImageJ program (NIH, Bethesda, MD).

### 2.6. Statistical analysis

Statistical analyses were performed using IBM SPSS Software version 20 (IBM Corp., NY). An independent *t*-test with two tails and unequal variances was used to determine the statistical significance of differences between the results of experimental and control groups. Descriptive data were summarized as mean ± standard error of mean.

## 3. Results and Discussion

### 3.1. Myocardial apoptosis is increased in C3 ^−/−^ mice after heart I/R

The current study used the mouse heart I/R model to investigate the effect of C3 on myocardial apoptosis. After 1 h ischemia and 24 hrs reperfusion, there were significantly more TUNEL-positive cells in hearts of C3^−/−^ mice compared with those of WT mice (3.4 ± 0.6 TUNEL positive nuclei/per mm^2^ vs. 1.2 ± 0.8 TUNEL positive nuclei/per mm^2^; P < 0.05) (Fig. 1. *i* to *iii*). To confirm this result, we analyzed *in situ* caspase-3 activation in the hearts of C3^−/−^ and WT mice after I/R. Immuno-staining of heart cryosections showed that post-I/R hearts of C3^−/−^ mice had more activated caspase-3 than those of WT mice (15.9 ± 1.8 positive area (μm^2^)/total area (mm^2^) vs. 1.1 ± 0.7 positive area (μm^2^)/total area (mm^2^); P < 0.05) (Fig.1. *iv* to *vi*). Thus, these results confirmed that myocardial apoptosis is increased in C3^−/−^ mice after 60 minutes of heart ischemia followed by 24 hrs of reperfusion. Taken together with our earlier report [13], these findings imply that in WT mice during myocardial I/R, C3 acts to promote necrosis and block apoptosis in heart.

**Fig. 1.**
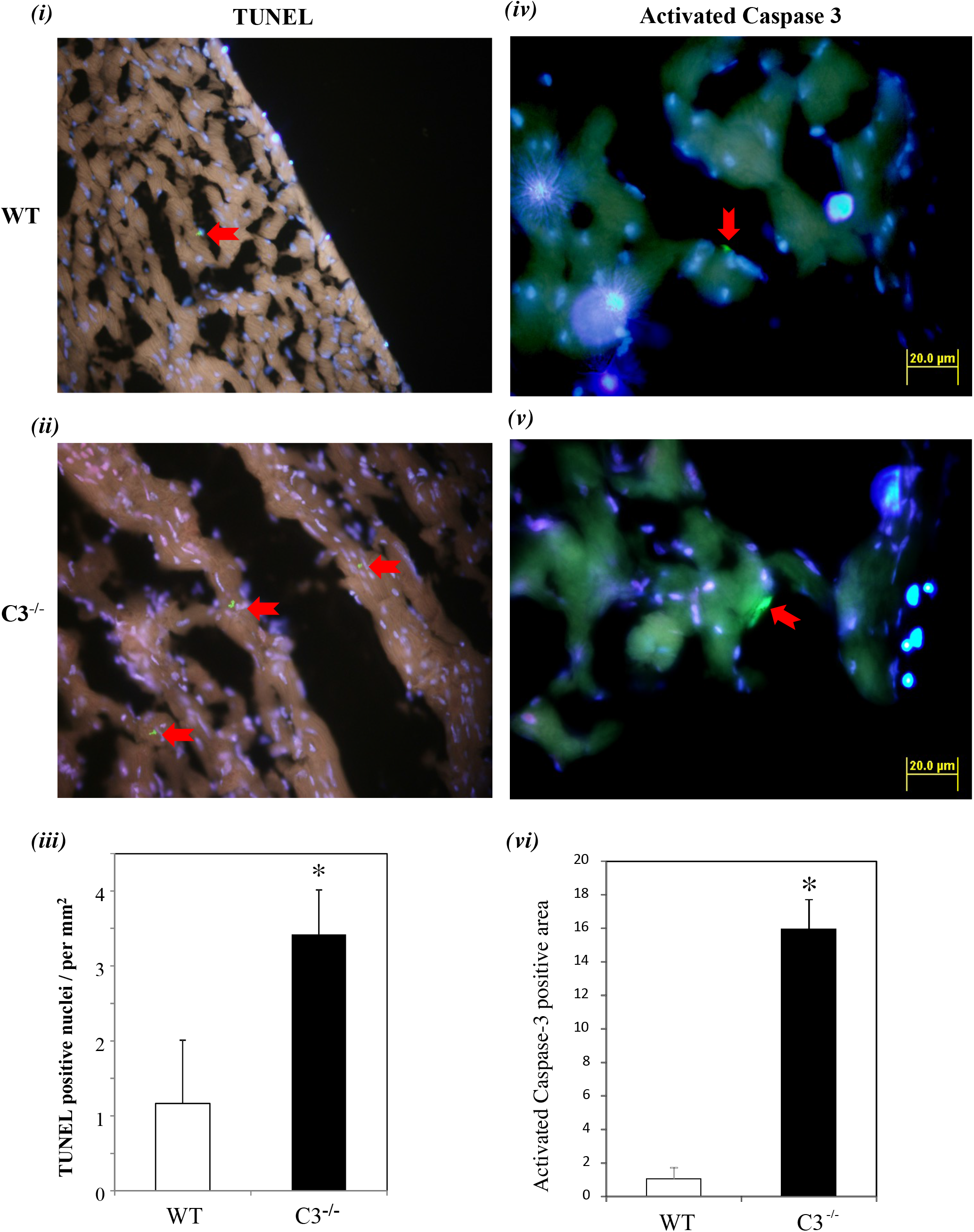
Apoptosis is increased in the hearts of C3^−/−^ mice after IR. WT and C3 ^−/−^ mice were subjected to 1 h myocardial ischemia and 24 hrs reperfusion. ***(i & ii)*** Apoptosis analyses by TUNEL assay. Cryosections of heart tissues were stained using a TACS 2TdT TUNEL kit according to the manufacturer’s directions. ***(iii)*** Bar graph: TUNEL-positive cells were quantified and expressed as numbers of positive nuclei/per mm^2^ of the cryosection and the average was obtained for the 3 mice/group. * *P*<0.05; error bars indicate SEM. ***(iv & v)*** Apoptosis analyses by activated caspase-3. Cryosections of hearts were stained for activated caspase-3 and area of positively stained cells were determined by fluorescent microscopy and analyzed using the ImageJ Program to quantify % Area of Positively Staining Cells compared to total Area of Tissue Section. ***(vi)*** Bar graph: bars indicate Area of Positively stained activated Caspase-3 / Total Area, in (μm^2^)/total area (mm^2^), for WT (n=5) and C3 ^−/−^ (n=5).

### 3.2. Identification of cytochrome c in a myocardial C3-complex following I/R using comparative proteomics

The increase of apoptosis in C3^−/−^ mice during I/R, led us to explore whether C3 might interact with factors related to programmed cell death. A comparative proteomic approach was used in our investigation. Our earlier study showed that in the mouse myocardial I/R model, C3 deposition becomes significant at 3 hrs reperfusion [12]. Similar C3 deposition was also reported by others in a rat model [21]. Thus, heart lysates were prepared from WT and C3^−/−^ mice after 1 hr ischemia/3 hrs reperfusion to capture the factor(s) interacting with C3 at the early phase. C3-containing protein complexes were isolated using anti-C3d conjugated Dynabeads (Life Technologies, CA). The complexes were analyzed by comparative proteomics (Fig. 2a). A total of 57 proteins were identified as being preferentially present in complexes with C3 in WT mice compared with C3^−/−^ mice (WT:C3^−/−^ ratio score >2, *P*<0.05)(Fig. 2b). Of these 57 proteins, only one protein, cytochrome *c*, is known to be directly involved in cell death (Supplementary Table 1). Pathway analysis predicted that cytochrome *c* is in a protein network that includes C3 (Supplementary Fig. 1). As far as we are aware of, there are no published reports for a direct interaction between the two. This experimental finding is the first evidence for a C3 complex containing cytochrome *c*.

**Fig. 2.**
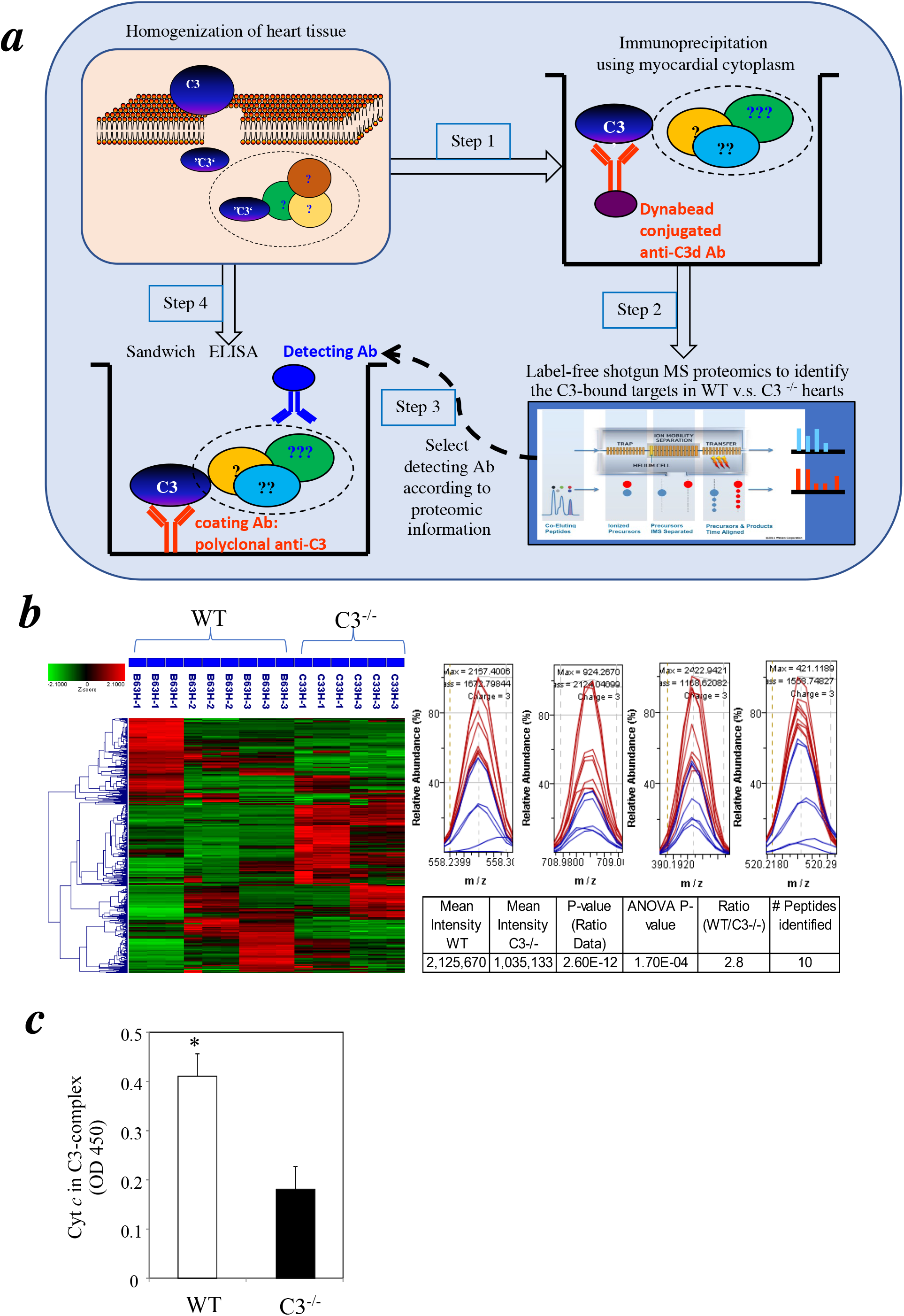
Comparative proteomics identified cytochrome *c* in C3-containing complexes formed during myocardial I/R. ***(a)*** Schematic diagram depicts the steps of experimental approaches. Step 1: hearts were harvested and homogenized from WT and C3^−/−^ mice which were subjected to 1 hr ischemia/3 hrs reperfusion. C3-containing complexes in heart lysates were immunoprecipitated with an anti-C3d Ab. Step 2: the C3-immunoprecipitated complexes were digested with trypsin and analyzed by label-free shot-gun LC-mass spectrometry. Step 3: Select detecting Ab according to proteomic information. Step 4: Sandwich ELISA to confirm the specific protein in C3-complex. ***(b)*** Left panel: Chromatograms were recorded for each biological replicate in the Resolution/Ion Mobility mode. Agglomerative hierarchical clusters of Z-score transformed intensity data were processed by the Elucidator program for all LC–MS chromatograms. Z-score coloration indicates protein abundance in WT compared with C3^−/−^ samples (red, higher abundance; green, lower abundance; black, equal abundance). Right panel histograms: Comparison of cytochrome *c* peptide signals of specific m/z ratios between WT and C3^−/−^ mice. Red and blue lines represent peptide signals from WT and C3^−/−^ mice, respectively. At bottom right is the summary table of the cytochrome *c* peptide signal comparisons of WT and C3^−/−^ mice. ***(c)*** Cytochrome *c* is present in a cytosolic C3-binding complex following 1h ischemia and 3 hrs of reperfusion. Myocardial cytosolic factions were isolated. C3-binding complexes in the myocardial cytosolic fractions were captured with a polyclonal anti-C3 Ab bound to the surface of microplates. An anti-cytochrome *c* Ab was used to detect cytochrome *c* in C3-binding complexes (n=3 mice/group). * *P*<0.05. Error bars indicate SEM.

It is relevant to note that cytochrome *c* is normally anchored to the inner mitochondrial membrane, but is released to the cytosol by various pro-apoptotic signals and induces apoptosis by activation of caspases [22,23]. The identification of cytochrome *c* in a C3-containing complex suggested that cytosol-located, activated C3 might interfere with the function of cytochrome *c* in apoptosis.

### 3.3. Detection of cytochrome c in a cytosolic C3-complex following I/R by immunoassay

To confirm the comparative proteomic results, we developed a sandwich ELISA to determine if cytochrome *c* and C3 interacted in the cytosol of post-I/R cardiomyocytes. Cytosolic fractions were isolated from the hearts of WT and C3^−/−^ mice after 3 hrs reperfusion and added to a microplate coated with a polyclonal anti-native C3 Ab. Cytochrome *c* was detected by an Ab against cytochrome *c*. There was significant binding of cytochrome *c* to C3 in WT mice compared with the background levels in C3^−/−^ mice (Fig. 2c).

Taken together, the results from proteomics and the cytosol ELISA assay indicate that C3 can interact directly with cytochrome *c* in the cytosol of cardiomyocytes during myocardial I/R.

### 3.4. Similar levels of cytochrome c in myocardial cytosol of C3^−/−^ and WT mice after I/R

It is well known that anti-apoptotic signaling molecules such as Bcl-2 and Bcl-xL prevent cytochrome *c* release from mitochondria and thus inhibit apoptosis [24]. Conversely, pro-apoptotic molecules, e.g. BAX and BH3-only proteins [25], enhance cytochrome *c* release and promote apoptosis. Our finding that C3^−/−^ mice have increased myocardial apoptosis after I/R compared with WT mice (Fig. 1) raised the question of whether C3^−/−^ mice have increased cytochrome *c* released into the cytosol after I/R.

To investigate this possibility, we isolated myocardial cytosolic fractions from C3^−/−^ and WT mice following I/R and compared the cytochrome *c* levels by Western blotting. There were no significant differences in the cytochrome *c* levels in C3^−/−^ and WT mice at each time point (Supplementary Fig. 2). Thus, the mitochondrial response to I/R stress, in terms of cytochrome *c* released, is similar in C3^−/−^ mice and WT mice.

Our results suggest that the increase in apoptosis in post-I/R C3^−/−^ hearts shown in Fig. 1 is not due to an increase in the amount of cytochrome *c* released into the cytosol, but to the genetic lack of C3 available for binding to cytochrome *c*. It is possible that binding of C3 to cytosolic cytochrome *c* in WT mice sequesters cytochrome *c*, thus reducing the number of cells which complete apoptosis. The result from our recent report indicates that necrosis is favored in WT mice expressing C3 [13]. Therefore, during I/R, based on our results, C3 minimizes apoptosis and promotes necrosis. These results raise the possibility of a new mechanism affecting cell death relevant to pathologic conditions such as ischemia: a circulating innate immune factor enters cardiomyocytes, interacts directly with intracellular factor(s), and influences the types of cell death that occur. Future studies on this line will provide more insights from basic science regarding regulation of I/R related myocardial cell death.

## Acknowledgements

The authors thank Drs. James Cottrell and David Wlody for their continued support.

