## Supplemental Table and Figures for "Complement C3 interacts with cytochrome *c* to influence myocardial apoptosis during heart ischemia/reperfusion"

**Supplementary Table 1. Comparative Proteomics Analyses of proteins in C3-binding complex**

| Primary Protein Name | Protein Description | P-value (Ratio Data) | ANOVA P-value | Ratio (WT/KO) | # Peptides |
|---|---|---|---|---|---|
| B2RQQ1_MOUSE | MCG133649, isoform CRA_a GN=Myh6; cardiac muscle, alpha; alpha-MHC | 0.01 | 0.03 | 2.2 | 125 |
| CO3_MOUSE | Complement C3, Alternative initiation; Cleavage on pair of basic residues; Complement alternate pathway; Complement pathway; Direct protein sequencing; Disulfide bond; Glycoprotein; Immunity; Inflammatory response; Innate immunity; Phosphoprotein; Secreted; Signal; Thioester bond | 1.8E-07 | 2.8E-03 | 2.3 | 47 |
| Q497E4_MOUSE | Actin alpha cardiac GN=Actc1 | 2.4E-20 | 1.9E-04 | 2.2 | 36 |
| MYL3_MOUSE | Myosin light chain 3 GN=Myl3 | 1.2E-09 | 4.3E-04 | 2.7 | 32 |
| ALBU_MOUSE | Serum albumin GN=Alb | 4.2E-07 | 6.9E-03 | 2.4 | 24 |
| Q545Y3_MOUSE | Tropomyosin 1, alpha, isoform CRA_l GN=Tpm1 | 6.2E-03 | 5.6E-03 | 18.5 | 22 |
| ACTN2_MOUSE | Alpha-actinin-2 GN=Actn2 | 3.9E-03 | 0.05 | 2.1 | 21 |
| B1AR69_MOUSE | Myosin, heavy polypeptide 13, skeletal muscle GN=Myh13 | 2.0E-06 | 0.04 | 2.7 | 19 |
| MYH3_MOUSE | Myosin-3 GN=Myh3 | 4.3E-03 | 0.08 | 2.3 | 19 |
| MYH7B_MOUSE | Myosin-7B GN=Myh7b | 4.4E-10 | 6.4E-04 | 2.6 | 13 |
| LDB3_MOUSE | Isoform Oracle 2 of LIM domain-binding protein 3 GN=Ldb3 | 5.2E-32 | 0.02 | 2.0 | 11 |
| CYC_MOUSE | Cytochrome c, somatic GN=Cycs | 2.6E-12 | 1.7E-04 | 2.8 | 10 |
| Q5SX41_MOUSE | Myosin, heavy polypeptide 2, skeletal muscle, adult GN=Myh2 | 2.1E-03 | 5.2E-04 | 5.8 | 8 |
| E9PZF0_MOUSE | Nucleoside diphosphate kinase GN=Nme2 | 0 | 0.30 | 2.6 | 8 |
| Q3UVB1_MOUSE | Myoglobin, isoform CRA_a GN=Mb | 1.2E-06 | 8.5E-03 | 2.0 | 7 |
| NDUS6_MOUSE | NADH dehydrogenase [ubiquinone] iron-sulfur protein 6, mitochondrial GN=Ndufs6 | 3.9E-03 | 0.18 | 2.0 | 7 |
| ECH1_MOUSE | Delta(3,5)-Delta(2,4)-dienoyl-CoA isomerase, mitochondrial GN=Ech1 | 9.8E-22 | 4.7E-03 | 3.2 | 6 |
| NDUAA_MOUSE | NADH dehydrogenase [ubiquinone] 1 alpha subcomplex subunit 10, mitochondrial GN=Ndufa10 | 5.3E-31 | 0.04 | 4.1 | 5 |
| B1ATS4_MOUSE | ATPase, Ca++ transporting, ubiquitous GN=Atp2a3 | 1.3E-35 | 1.4E-06 | 3.2 | 5 |
| K2C75_MOUSE | Keratin, type II cytoskeletal 75 GN=Krt75 | 4.2E-33 | 8.0E-07 | 8.7 | 4 |
| B1ASG5_MOUSE | Ubiquinol-cytochrome c reductase hinge protein GN=Uqcrh | 1.5E-10 | 7.1E-07 | 7.7 | 4 |
| Q497F1_MOUSE | Troponin I, cardiac 3 GN=Tnni3 | 9.8E-05 | 5.9E-04 | 4.4 | 4 |
| E9PYX4_MOUSE | Glyceraldehyde-3-phosphate dehydrogenase | 4.1E-08 | 2.8E-03 | 3.6 | 4 |
| K2C1_MOUSE | Keratin, type II cytoskeletal 1 GN=Krt1 | 5.3E-08 | 3.6E-04 | 3.3 | 4 |
| K22O_MOUSE | Keratin, type II cytoskeletal 2 oral GN=Krt76 | 2.1E-03 | 0.07 | 3.0 | 4 |

| | | | | | |
|---|---|---|---|---|---|
| TBA4A_MOUSE | Tubulin alpha-4A chain GN=Tuba4a | 1.9E-11 | 3.8E-03 | 2.5 | 4 |
| E9Q264_MOUSE | Uncharacterized protein GN=Myh15 | 0.10 | 0.44 | 2.1 | 4 |
| SODC_MOUSE | Superoxide dismutase [Cu-Zn] GN=Sod1 | 1.4E-05 | 2.8E-04 | 4.7 | 3 |
| E9QPE7_MOUSE | Uncharacterized protein GN=Myh11 | 0.08 | 0.82 | 3.6 | 3 |
| Q3UH59_MOUSE | Myosin, heavy polypeptide 10, non-muscle GN=Myh10 | 0.06 | 0.37 | 3.5 | 3 |
| ANR53_MOUSE | Ankyrin repeat domain-containing protein 53 | 4.3E-06 | 0.01 | 3.2 | 3 |
| A2A6Q8_MOUSE | Myosin, light polypeptide 4 (Fragment) GN=Myl4 | 4.2E-12 | 0.17 | 3.1 | 3 |
| Q543D7_MOUSE | Four and a half LIM domains 2, isoform CRA_a GN=Fhl2 | 4.6E-03 | 0.04 | 2.8 | 3 |
| Q6GT24_MOUSE | Peroxiredoxin 6 GN=Prdx6 | 5.7E-10 | 4.8E-03 | 2.5 | 3 |
| K6PL_MOUSE | 6-phosphofructokinase, liver type GN=Pfkl | 1.9E-06 | 0.01 | 2.4 | 3 |
| E9Q607_MOUSE | Uncharacterized protein GN=Actg1 | 3.3E-04 | 0.23 | 2.3 | 3 |
| H4_MOUSE | Histone H4 GN=Hist1h4a | 4.2E-07 | 4.5E-03 | 2.2 | 3 |
| B2RTK3_MOUSE | Histone H2B GN=Hist1h2bm | 8.2E-06 | 0.04 | 2.2 | 3 |
| PLIN4_MOUSE | Perilipin-4 GN=Plin4 | 2.0E-08 | 8.7E-05 | 2.0 | 3 |
| K6PF_MOUSE | 6-phosphofructokinase, muscle type GN=Pfkm | 4.9E-04 | 1.6E-03 | 9.5 | 2 |
| B2RXX9_MOUSE | Myosin, heavy polypeptide 7, cardiac muscle, beta GN=Myh7 | 1.4E-15 | 2.6E-03 | 8.2 | 2 |
| E9QLL7_MOUSE | Uncharacterized protein GN=Itga7 | 3.8E-09 | 7.9E-05 | 7.9 | 2 |
| Q546G4_MOUSE | Albumin 1 GN=Alb | 2.5E-06 | 1.1E-04 | 7.7 | 2 |
| RT36_MOUSE | 28S ribosomal protein S36, mitochondrial GN=Mrps36 | 1.7E-08 | 1.0E-04 | 7.1 | 2 |
| F8WIG3_MOUSE | Uncharacterized protein GN=Ninl | 3.3E-05 | 6.0E-03 | 6.6 | 2 |
| B2RRX1_MOUSE | Actin, beta GN=Actb | 1.0E-05 | 4.1E-03 | 6.6 | 2 |
| LDHC_MOUSE | L-lactate dehydrogenase C chain GN=Ldhc | 5.8E-12 | 1.1E-04 | 4.6 | 2 |
| NDUA7_MOUSE | NADH dehydrogenase [ubiquinone] 1 alpha subcomplex subunit 7 GN=Ndufa7 | 1.7E-24 | 3.4E-05 | 4.5 | 2 |
| E9QAS7_MOUSE | Uncharacterized protein GN=Inpp5a | 5.6E-03 | 0.24 | 4.2 | 2 |
| A1AT5_MOUSE | Alpha-1-antitrypsin 1-5 GN=Serpina1e | 0.27 | 0.78 | 3.5 | 2 |
| IGH1M_MOUSE | Ig gamma-1 chain C region, membrane-bound form GN=Ighg1 | 5.4E-18 | 0.04 | 3.4 | 2 |
| D3Z0Z9_MOUSE | Glyceraldehyde-3-phosphate dehydrogenase GN=Gm5069 | 2.4E-30 | 4.9E-04 | 3.3 | 2 |
| QCR7_MOUSE | Cytochrome b-c1 complex subunit 7 GN=Uqcrb | 8.5E-15 | 8.5E-05 | 3.1 | 2 |
| A2AIR5_MOUSE | Glutamate receptor ionotropic, NMDA3A GN=Grin3a | 2.3E-05 | 1.9E-03 | 3.0 | 2 |
| A2AEW8_MOUSE | GRIP1 associated protein 1 GN=Gripap1 (neuron-specific guanine nucleotide exchange factor for the Ras family of small G proteins (RasGEF) and is associated with the GRIP/AMPA receptor complex in brain) | 3.2E-25 | 6.4E-04 | 2.9 | 2 |
| E9Q1N7_MOUSE | Uncharacterized protein | 0.01 | 0.13 | 2.7 | 2 |

| | | | | | |
|---|---|---|---|---|---|
| COG6_MOUSE | Conserved oligomeric Golgi complex subunit 6 GN=Cog6 | 3.2E-03 | 9.2E-03 | 2.7 | 2 |
| E9Q452_MOUSE | Uncharacterized protein GN=Tpm1 | 2.6E-03 | 0.01 | 2.2 | 2 |
| B1AXW5_MOUSE | Peroxiredoxin 1 (Fragment) GN=Prdx1 | 1.2E-06 | 0.03 | 2.2 | 2 |

**Supplementary Figure 1**

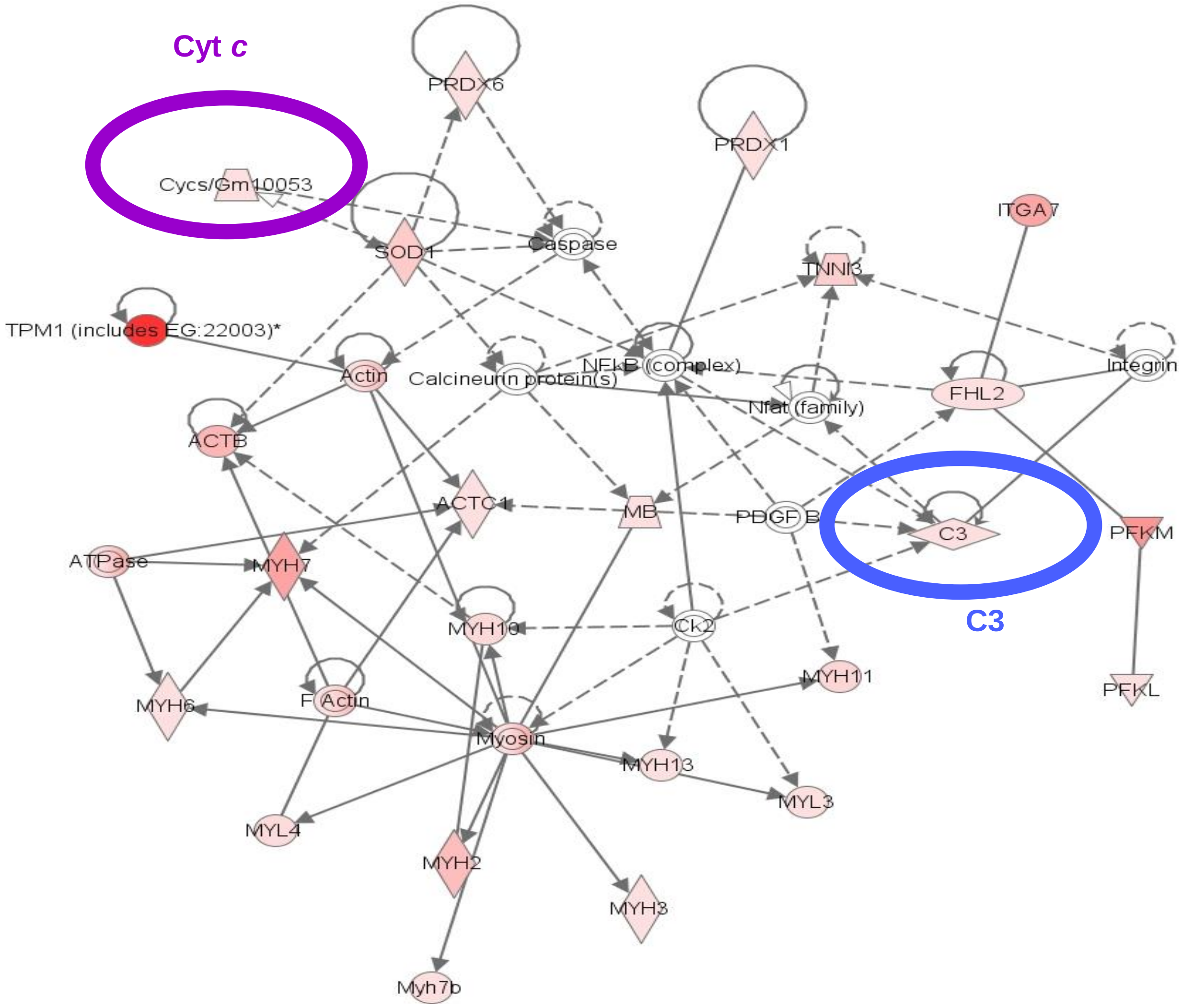


**Supplementary Fig. 1. Pathway analysis of protein networks associated with the C3**. The network was generated by Ingenuity pathway analysis (IPA) software using the list of differentially expressed 57 proteins identified by proteomics. The yellow circle on the left highlights cytochrome *c*, that on the right highlights C3.

### Supplementary Figure 2

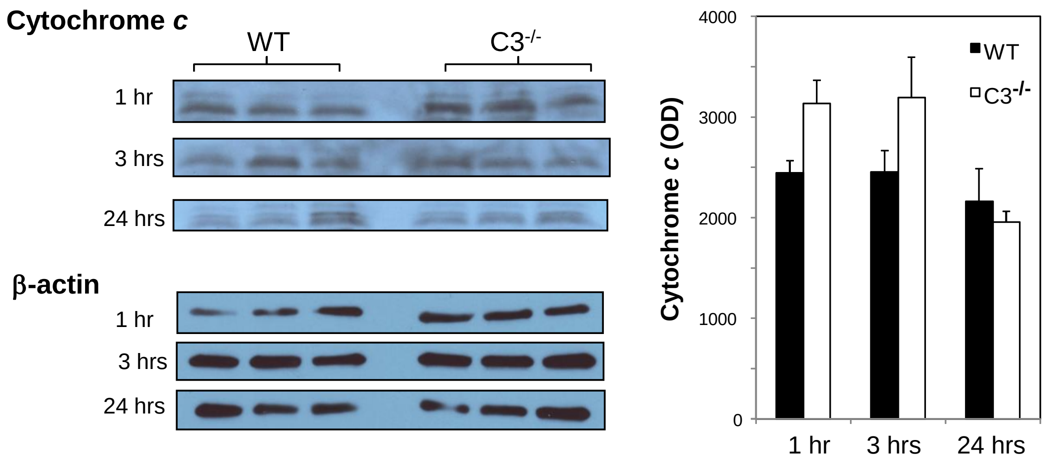


**Supplementary Fig. 2. Cytochrome *c* levels in $C3^{-/-}$ and WT myocardial cytosols after I/R.** Cytosolic factions were isolated and the proteins separated by SDS-PAGE. Western blotting was carried out using an anti-cytochrome *c* Ab and ECL detection. β-actin, detected with an anti-β-actin Ab, acted as a control for cytosol volume added/well. The left panels are blots of cytochrome *c* and β-actin. Each lane represents an individual mouse. The right chart shows the results of densitometric analysis of the cytochrome *c* bands averaged for the 3 mice. $P > 0.05$ for $C3^{-/-}$ and WT mice results.
